# The peptide sensor motif stabilizes the outward-facing conformation of TmrAB

**DOI:** 10.1101/2020.01.12.903617

**Authors:** Cinthia R. Millan, Martina Francis, Valery F. Thompson, Tarjani M. Thaker, Thomas M. Tomasiak

## Abstract

The ATP binding cassette (ABC) family of transporters move diverse small molecules across membranes in nearly all organisms. Transport activity requires conformational switching between inward-facing and outward-facing states driven by ATP-dependent dimerization of two nucleotide binding domains (NBDs). The allosteric mechanism that connects ATP binding and hydrolysis in the NBDs to conformational changes in a substrate binding site in the transmembrane domains (TMDs) presents an unresolved question. Here we use sequence coevolution analyses together with biochemical characterization to investigate the role of a highly conserved motif called the peptide sensor in coordinating domain rearrangements in the heterodimeric peptide exporter from *Thermus thermophilus*, TmrAB. Mutations in the peptide sensor motif alter ATP hydrolysis rates as well as substrate release. Disulfide crosslinking, evolutionary trace, and evolutionary coupling analysis reveal that these effects likely destabilize a network between the peptide sensor motif and the Q-loop and X-loop, two known allosteric elements in the NBDs. We further find that disruption of this network in TmrA versus TmrB has different functional consequences, hinting at an intrinsic asymmetry in heterodimeric ABC transporters extending beyond that of the NBDs. These results support a mechanism in which the peptide sensor motifs help coordinate the transition of TmrAB to an outward open conformation, and each half of the transporter likely plays a different role in the conformational cycle of TmrAB.

## INTRODUCTION

ATP binding cassette (ABC) transporters constitute a vast protein superfamily found in every domain of life. They consume ATP to transport biosynthetic components such as lipids or toxic molecules including drugs, toxins, and peptides across membranes [1,2]. Their dysfunction is genetically linked to many human diseases including cystic fibrosis, Stargardt disease, Tangier’s disease, and multiple dyslipidemias [3]. Because of their medical relevance, the transport mechanism has long been a focal point of ABC transporter research, especially of exporters which predominate in humans. Structural snapshots of isolated states reveal that transport in both ABC exporters and importers is accompanied by large rearrangements between a pair of nucleotide binding domains (NBDs) that bind and hydrolyze ATP and a pair of transmembrane domains (TMDs) that bind and transport substrate.

The domain rearrangements that underlie the transport cycle in ABC transporters progresses through three general steps: 1) ATP-driven NBD dimerization, 2) conformational change from a high affinity inward-facing (IF) state to a low affinity outward-facing (OF) state, and 3) ATP hydrolysis to reset the transport cycle (**Figure 1A**). Alternative models that build on this general mechanism have been proposed by biochemical and genetic studies [4–6], high resolution structures determined by x-ray crystallography and cryogenic electron microscopy (cryo-EM) in different states and conformations [7–12], as well as distance measurements by electron paramagnetic resonance (EPR) [13,14]. The most widely recognized models proposed can be classified into two primary groups based on mechanistic commonalities [15]. Models in the first category suggest a complete separation of NBDs occurs after each cycle and includes the Switch model [16], the Tweezers-like model [17], and processive clamp model [18]. In contrast, the second category proposes a continuous contact of at least one NBD during the transport cycle and is comprised of the nucleotide occlusion model [19,20], the alternating catalysis model [21] and the constant contact model [22]. These models serve to define the mechanism behind substrate release and the precise role of ATP binding versus hydrolysis in transporter function. Together, their findings suggest transport cycles can proceed via symmetric or asymmetric NBD dimerization - the latter being associated with asymmetric transporter architectures. Nevertheless, all of them strongly support the role of ATP binding to bridge two NBDs and initiate a conformational change in the TMD during transport.

**Figure 1.**
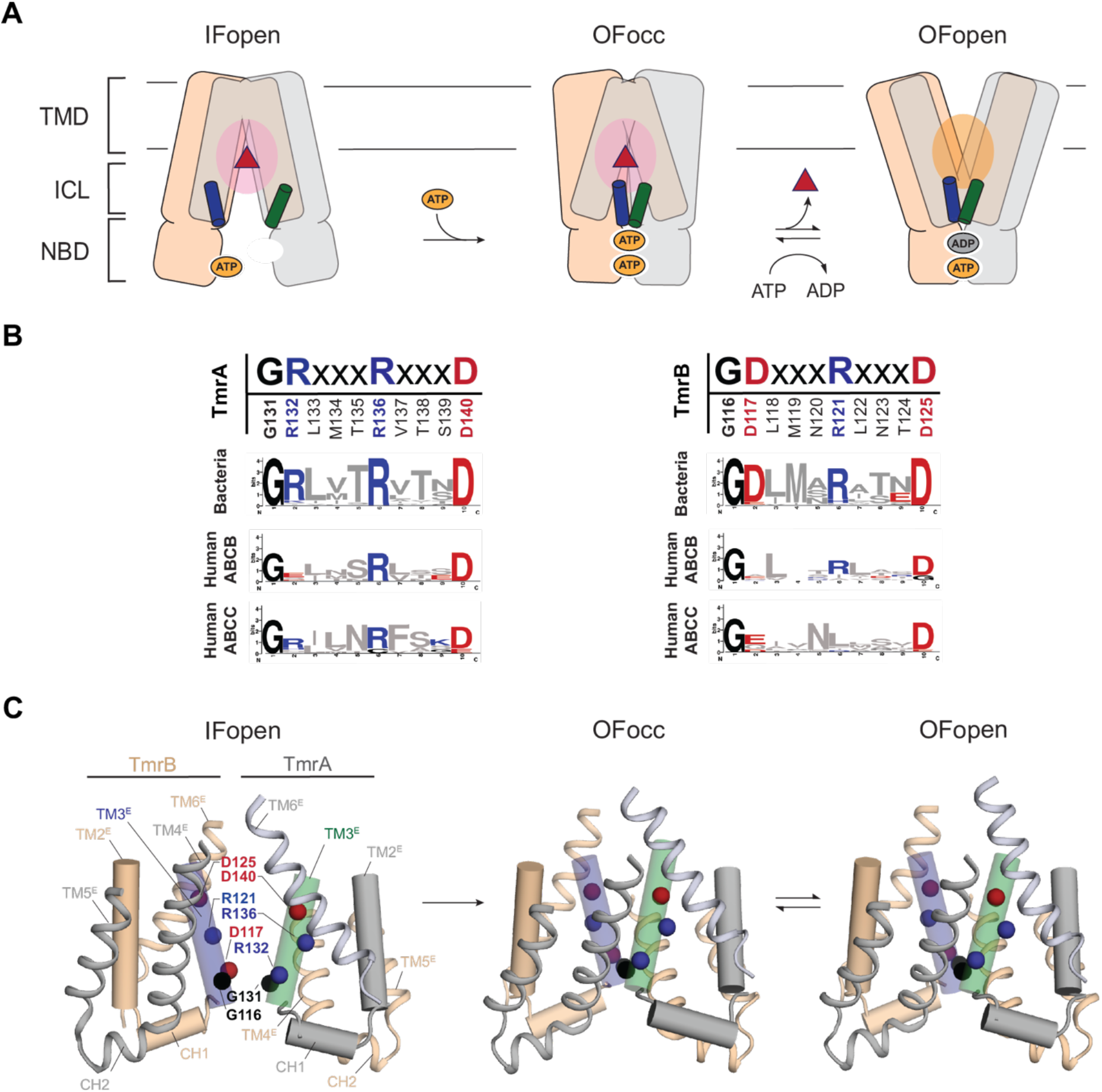
The conserved Peptide Sensor Motif in intracellular loop 1 creates critical connections upon transition to the outward open state during the transport cycle of TmrAB. **A**) Cartoon overview of the inward-facing open (IFopen) to outward-facing occluded (OFocc) and outward-facing open (OFopen) transitions of TmrAB highlighting relative orientations of the transmembrane domains (TMD), intracellular loops (ICL), and nucleotide binding domains (NBD) from TmrA (*grey*/*green*) and TmrB (*orange/blue*) at the start and end of the proposed transport cycle. The helical region containing the Peptide Sensor Motif (PSM) in intracellular loop 1 is shown in *green* for TmrA and in *blue* for TmrB. **B**) PSM sequence in the canonical Walker B containing TmrA chain (left) and non-canonical Walker B motif containing TmrB chain (right), with PSM consensus sequences for related ABC transporter families for each chain shown below. **C**) Cartoon representation of the local rearrangements in the ICLs of TmrA and TmrB in the IFopen (left, PDBID: 6raf [10]), OFocc (middle, PDBID: 6rak [10]), and OFopen (right PDBID: 6raj [10]) states. Intracellular loop 1 in TmrA (*grey/green*) and TmrB (*orange/blue*) comprised of cytoplasmic extensions from transmembrane helices 2 (TM2^E^) and 3 (TM3^E^) connected by coupling helix 1 (CH1) are shown as cylindrical helices. The transmembrane helix 6 extension (TM6^E^) linking the TMD and NBD and the intracellular loop 2 comprised of extensions from transmembrane helices 4 (TM4^E^) and 5 (TM5^E^) connected by coupling helix 2 (CH2) are shown as cartoon representations colored *grey* in TmrA and *orange* in TmrB. Conserved residues of the PSM on the TM3^E^ helix are shown as spheres colored the same as in (**B**).

A key element for conformational coupling is allosteric communication between ATP binding sites and the substrate binding sites. As an example, substrate binding often stimulates or, in some cases, inhibits ATPase activity [23]. Investigations into their allosteric relationship have focused on two motifs in each NBD, the Q-loop and X-loop [24–26] (**Supplemental Figure 1**). Both motifs are found on loops that bridge the NBDs to the TMDs and are highly conserved, with the X-loop found only in ABC exporters [27]. Mutations in the Q-loop of the drug transporter human P-glycoprotein (PgP) result in defects in the ability to transition to outward open and a decoupling of ATPase activity and substrate transport [28]. Similarly, X-loop mutations in ABCB4 found in progressive familial intrahepatic cholestasis type-3 (PFIC-3), a liver disease involving aberrant bile formation, result in uncoupling of substrate binding and ATPase activity [29]. Both the X- and Q-loop interact with the TMD through helical extensions of the transmembrane forming intracellular loops (ICL) bridging these NBD motifs to the substrate binding site in the TMD (**Figure 1A**) [24].

Comparatively less emphasis has been placed on the ICLs compared to the X- and Q-loops despite being poised to provide a mechanical link between the ATP and substrate binding sites. Sequence similarity is generally lower in the TMDs than in the highly conserved NBDs, likely owing to the variability of transported substrates. This variability makes identifying conserved allosteric pathways difficult, but several recent investigations support a sequence specific role of intracellular loop 1 (ICL-1) in the transport cycle. Mutations in a conserved patch of charged residues in ICL-1 of the bacterial lipid A transporter, MsbA, destabilize the outward facing state during its transport cycle [30]. Furthermore, cysteine crosslinking in the human T-cell Antigen Presenting complex, TAP1/TAP2, peptide transporter showed that residues defined as the “peptide sensor” motif (abbreviated as the PSM here) decreased transport and altered interactions throughout the transport cycle [2]. The peptide sensor sequence (Gly(Arg/Asp)x_3_Argx_3_Asp) is conserved in the evolutionarily related homolog of TAP1/TAP2, TmrAB from *Thermus thermophilus* (**Figure 1B**). TmrAB is a peptide transporter with similar substrate specificity to TAP1/TAP2, and notably, can functionally replace TAP1/TAP2 in cell-based experiments [31]. The conservation in sequence and function by a bacterial ancestor of this class of proteins suggests an important role of this understudied motif in the regulation of ABC transporters, and further investigations may provide insights into their conformational cycle.

TmrAB is one of the best characterized ABC transporters to date [6,10,31–33]. A recent study reported cryo-EM structures of 6 different states of TmrAB [10]. This complete view of its conformational landscape allowed us to identify the PSM as a potential regulatory motif in a heterodimeric asymmetric ABC transporter. In this class, one NBD lacks the ability to hydrolyze ATP due to a non-canonical walker B motif (ϕϕϕϕDD instead of the canonical ϕϕϕϕDE – where ϕϕϕϕ is any hydrophobic residue), a feature shared with many medically relevant human transporters including TAP1/TAP2, the Cystic Fibrosis Transmembrane Regulator (CFTR), and Multidrug Resistance Protein (MRP) proteins. In TmrAB, the canonical walker B motif is present in the TmrA NBD and the non-canonical walker B motif is found in the TmrB NBD. We observed that interactions formed by the peptide sensors in the structures of both TmrA and TmrB are substantially altered as the protein proceeds through the proposed transport cycle (**Figure 1C**). Upon transition to the outward open state, a new helical bundle is formed between the PSMs of both chains and cytoplasmic extensions of the TMDs. The symmetry of the tightly assembled ICLs is noteworthy especially in light of the functional asymmetry between TmrA and TmrB and raises the question whether there is a role for peptide sensor residues in facilitating ATP-dependent conformational coupling in TmrAB.

Here, we further investigate the hypothesis that the peptide sensor region acts as a physical couple in TmrAB conformational changes upon ATP binding. Our analysis builds on previous characterization of the peptide sensor region from TAP1/TAP2 [2,24] and further defines the Gly(Arg/Asp)x_3_Argx_3_Glu as a peptide sensor motif (PSM). We show consensus in this sequence across ABC exporters using large scale evolutionary analysis with TmrAB and representative members of every class of human exporter that display a similar fold (B, C, and D) and of their bacterial homologs. In the PSM, the charged residues are highly conserved, and the glycine residue is almost absolutely conserved. Biochemically, we observe asymmetric disruption of the ATPase catalytic cycle when either PSM is disrupted, suggesting different roles of the PSM in the catalytically active and inactive chain of the transporter. Furthermore, we find that the highly conserved charged residues specifically are critical for stabilizing the outward open conformation necessary for export. Together, our results shed insights on a potential mechanism for stabilizing the formation of structural intermediates in the conformational transitions underlying ABC transporter activity and helping propagate allosteric information between ATP binding and substrate binding sites.

## RESULTS

### The peptide sensor motif forms part of an evolutionarily constrained network connecting nucleotide binding to the transmembrane region

To assess the extent to which the peptide sensor motif (PSM) within intracellular loop 1 (ICL-1) may be important for allosteric nucleotide coupling in asymmetric ABC transporters, we first analyzed its conservation across evolution (**Figure 1B, Supplemental Data 1, Supplemental Figure 1**). Sequences of bacteria and homologs in humans (the B, C, and D families) were selected and aligned to TmrA and TmrB separately to highlight potential relationships between PSM conservation and catalytic competency of the NBD. In a large scale comparison of 615 TmrA-like (**Figure 1B**,left) and 567 TmrB-like (**Figure 1B**,right) bacterial ABC transporter sequences identified using only the transmembrane regions of TmrAB to limit folds dissimilar to TmrAB, the Gly(Arg/Asp)x_3_Argx_3_Asp sequence in TmrAB can be identified as one of the most conserved sequences in the transmembrane region. Remarkably, the glycine in position 1 was 100% conserved in bacterial transporters similar to TmrA (G131^TmrA^) and in TmrB (G116^TmrB^), as well as in human homologs (**Figure 1B, Supplemental Data 1, Supplemental Figure 1**). Similarly, high identity or near identity was found in the charged residues in both TmrA-like (R132 -~90% R or K, R135-100% R or K, D140 – 100%) and TmrB-like (D117 – 97% D or E, R120 – 83% with R or K, D125 – 100%) sequences of ABC exporters in bacteria. Nonetheless, in all sequences a glycine residue precedes three charged residues that follow an *i* + 4 helical pattern (i.e. are spaced by 3 residues). This putative consensus motif connects the coupling helix-1 (CH1) positioned at the surface of the NBD to transmembrane helix 3 (TM3) in the TMD in TmrAB [2,24] (**Figure 1C**).

### Evolutionary contact predictions show the PSM connects NBDs to the transmembrane in the outward-occluded and outward-facing state of TmrAB

The scale of our sequence alignments allowed us to take a closer look at the functional relevance of the PSM in ABC exporters through sequence coevolution analysis. Many predictive techniques for classifying the structural and functional properties of specific sites through multi-sequence analysis have emerged in recent years [34]. The Evolutionary Trace method is one such approach that is particularly powerful because it can infer likely functionally important features from phylogenetic relations and information content through evolution of a family [35]. Using the Universal Evolutionary Trace method [36], residues of TmrA and TmrB were assigned a coverage score correlating with functional importance and in which residues with a lower score (higher inverse score) are considered more functionally important. As expected, residues in multiple known motifs in the NBDs ranked in the top 5% (above 95% rank) (**Figure 2A**). This tier surprisingly also included several sites in the TmrAB ICLs (4 in ILC-1 and 3 in ICL-2 of TmrA; 2 in ICL-1 and 1 in ICL-2 of TmrB). These include residues in position 1 (G131), 6 (R136), and 10 (D140) in TmrA and positions 1 (G116) and 10 (D125) in TmrB of their conserved PSM motifs.

**Figure 2.**
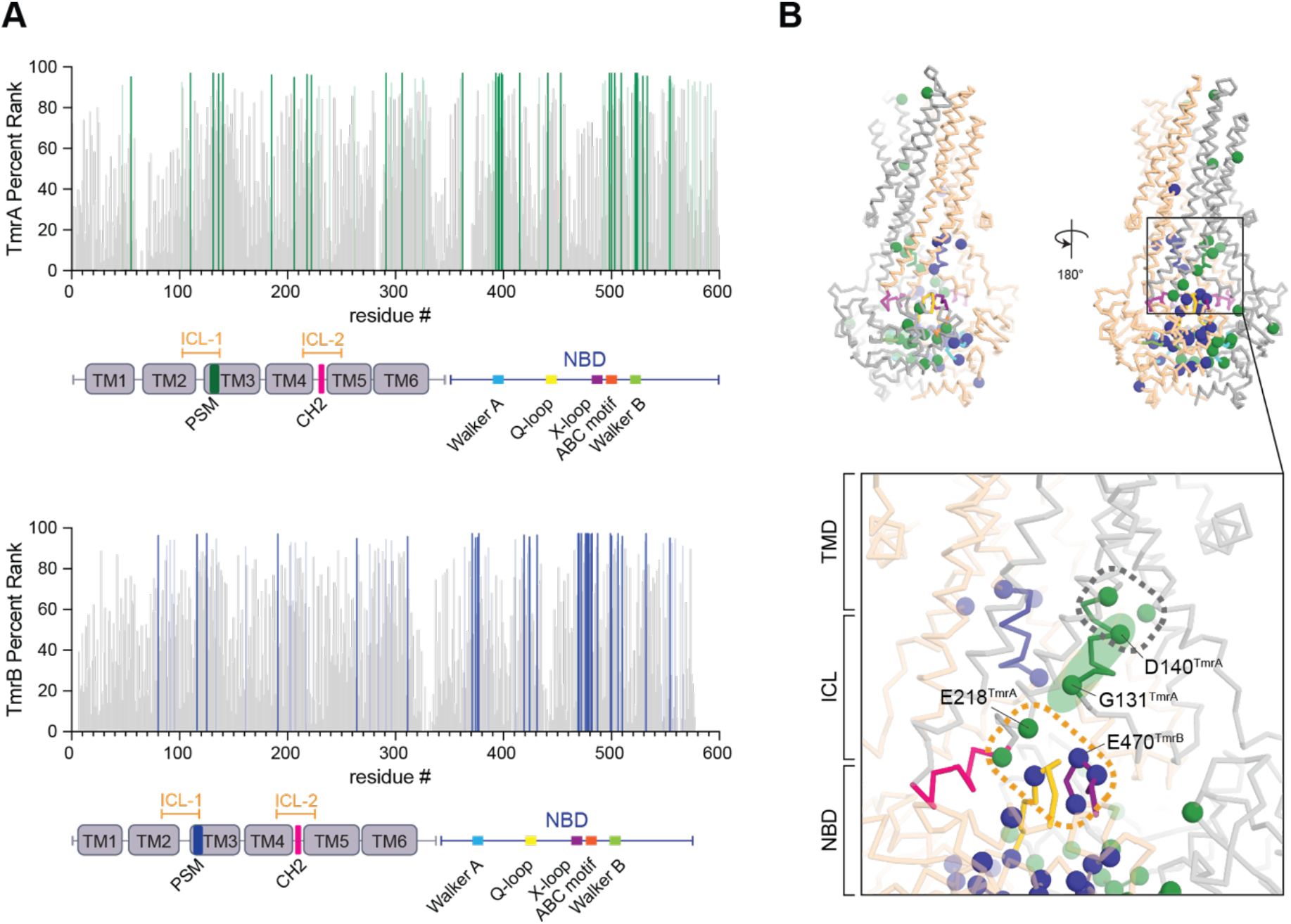
Evolutionary Trace highlights PSM-mediated networks bridging the TmrAB NBD and TMD as a potential route for conformational coupling. **A)** Real valued evolutionary trace scores using the Universal Evolutionary Trace server [64] plotted by residue with top scoring residues highlighted for TmrA (*dark green* – top 5%, *light green* – top 10%) and TmrB (*dark blue* - top 5%, *light blue* – top 10%). **B**) Top 5% of TmrA (*green spheres* on *grey*) and TmrB (*blue* spheres on *orange*) mapped onto the OFopen TmrAB structure (PDBID: 6raj [10]), with the inset highlighting the continuous network bridging the NBD to the TMD. Residues of the TMD in TmrA are highlighted by a *grey* dashed box, the PSM of TmrA is highlighted by a *green* oval, and the NBD in TmrB is highlighted by an *orange* dashed box.

Further analysis of all sites above 95% rank in the context of the TmrAB conformational states reveals a distinct continuous network that extends from the nucleotide binding sites at the NBD dimer interface, upwards along the peptide sensor in the TmrA ICL-1, and into the TMD in the outward-open TmrAB structure (PDB ID: 6raj [10]; **Figure 2B**). This network is anchored by G131^TmrA^ at the NBD/ICL-2 and ICL-1 interface and by D140^TmrA^ at the ICL-1/TMD interface. Interestingly, this putative route is not conserved on the TmrB face of the heterodimer when considering only the top 5% of sites, and lacks the cluster that forms the NBD connection to the TmrB ICL-1 as observed in TmrA, highlighting one possible route for conformational coupling between the ATP binding and substrate binding sites.

To better understand the structural basis for how the peptide sensor motif (PSM) may regulate the NBD and the TMD, we investigated evolutionary sequence covariations in TmrAB using a second evolutionary mapping approach and the EVCouplings software suite (**Figure 3**) [37]. Evolutionary coupling analysis provides direct insights into 3-dimensional contacts through the use of large scale sequence alignments (typically several hundred to several thousand sequences) and accounts for global coupling of sequences that are better able to filter indirect contacts that arise in methods such as Evolutionary Trace [35]. The success of these methods has been best displayed in the success of using these contacts for *de novo* structure prediction [38]. The EV couplings analysis of TmrA (**Figure 3A**) and TmrB (**Figure 3B**) shows extensive contacts through the PSM as is noted in the sequence alignments and Evolutionary Trace results above. Contacts predicted by the structures of TmrAB, notably outward-facing and outward-occluded, are captured by EV couplings to a high degree and include the aspartate in the 10^th^ position of the PSM. Notably, only coevolving positions within spatial proximity observed in known TmrAB structures are found, suggesting that the conformational changes between them may be important to explain allostery.

**Figure 3.**
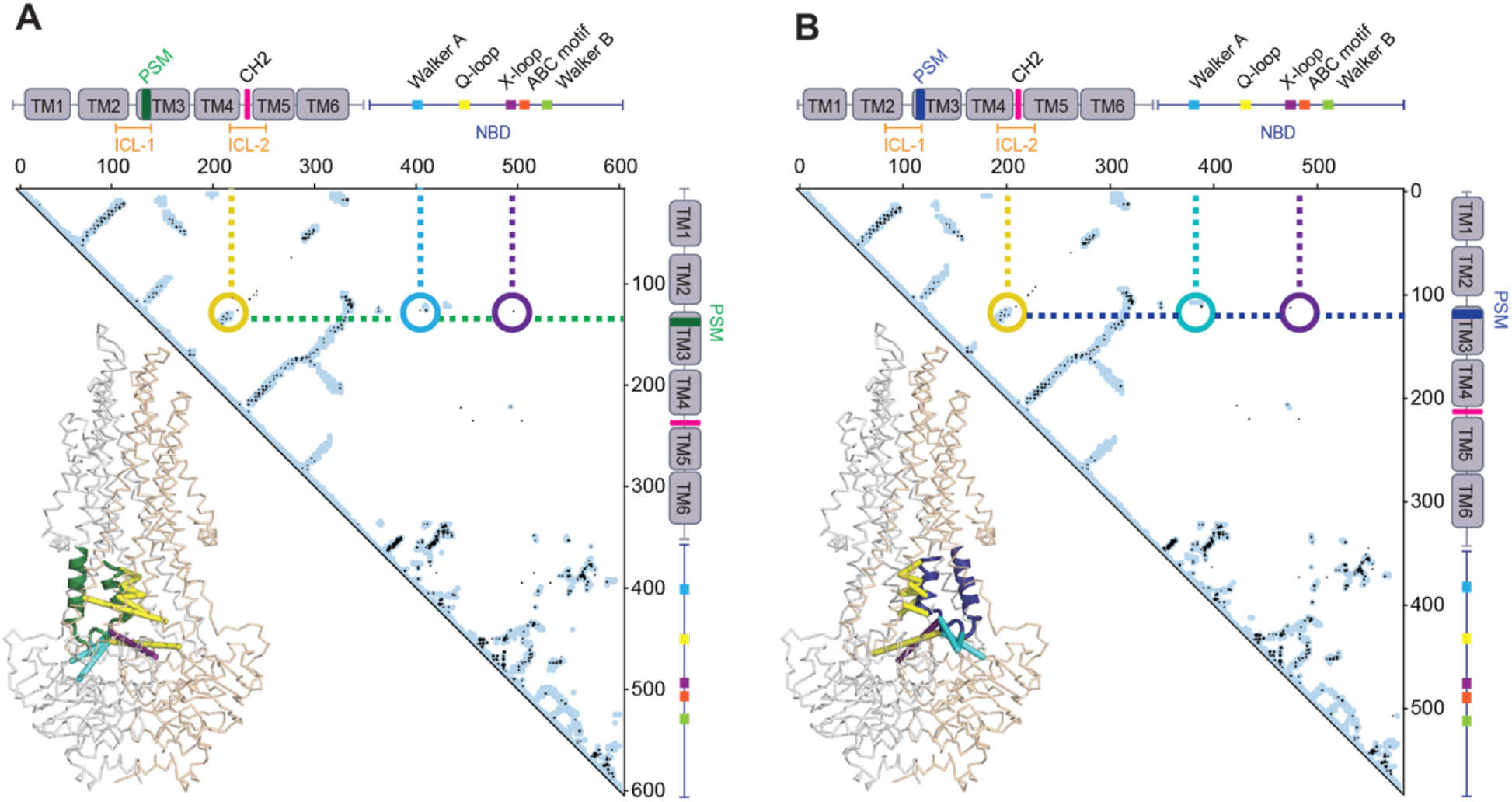
EVcouplings analyses show a network of co-evolving residue pairs between the PSM and the ATP binding sites in the outward-facing TmrAB state. Contact maps displaying intra-evolutionary couplings with an EVcouplings probability ≥0.99 (black dots) in **A**) TmrA and **B**) TmrB. Spatially close contacts predicted using structural information on TmrAB are highlighted in *light blue* in both. Dashed lines in the contact maps highlight the conserved evolutionary connections (with a probability > 0.99) between the PSM in TmrA (*green*) and TmrB (*blue*) to residues in intracellular loop 2 (ICL-2; *yellow*), and in proximity to the Walker A motif (*cyan*), the ABC motif, and the X-loop motif (*purple*) in the NBD. These interactions are mapped as distances onto a cartoon representation of the OFopen structure of TmrAB (PDBID: 6raj [10]) in their respective chains and colored the same as in the contact map.

Closer inspection of the EV couplings show important interactions that are only present in the outward open state that may be important for allostery. The most important connections arise between the PSM and the ATP binding site in both TmrA and TmrB through residues of the X-loop and the Walker A motif (**Figure 3**). We compared the number of connections in ICL-1 and ICL-2 and detected higher conserved co-evolving residues pairs for ICL-1 over ICL-2 (4.6x). Important connections between the NBD and ICL-1, but not with ICL-2, were identified. This suggests a potential role of ICL-1-mediated coupling of the transmembrane domain and NBDs.

### TmrAB PSM mutants exhibit WT like thermostability

To test the role of specific PSM residues in TmrAB activation, we generated TmrAB PSM consensus motif mutations in TmrA (G131A, R132A, R136A, and D140A) and in TmrB (G116A, D117A, R121A, and D125A). Intrinsic tryptophan fluorescence denaturation experiments performed on a Nanotemper Tycho show that all of the variants were properly folded with no significant changes in melting temperatures (>90°C) compared to wild-type (WT) TmrAB (**Figure 4**) with similar denaturation profiles. Furthermore, samples were inspected for aggregation using negative stain electron microscopy, which did not reveal perturbations to the overall architecture (**Supplemental Figure 2**). These results suggest that the PSM mutants do not interfere with stability of the resting folding state of TmrAB.

**Figure 4.**
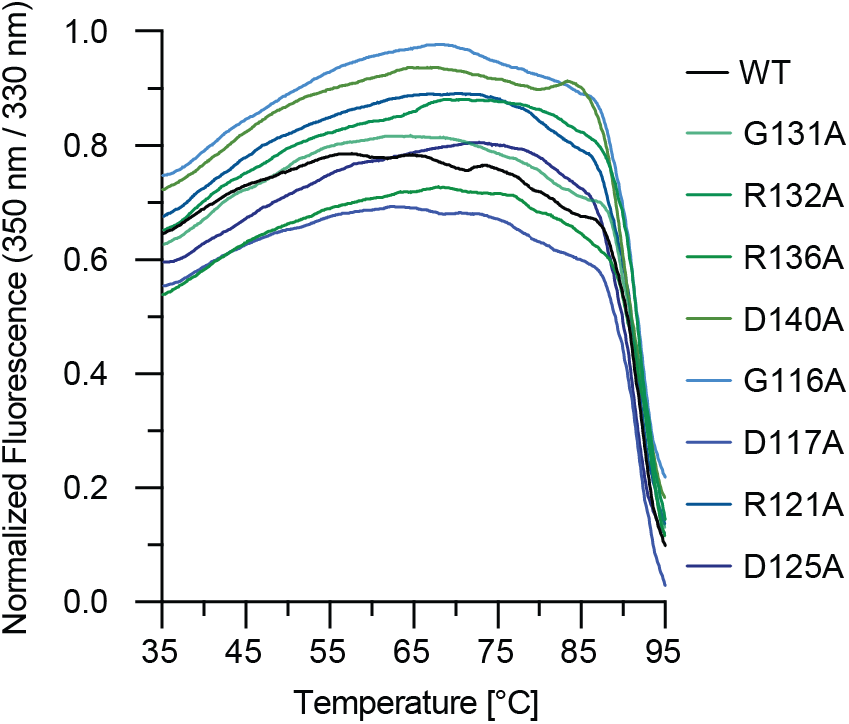
PSM mutations do not affect protein stability in TmrAB. Thermostability of WT and TmrAB peptide sensor mutants measured by intrinsic tryptophan fluorescence performed for purified samples in buffer containing 20 mM HEPES pH 7.0, 150 mM NaCl and 0.05% ß-DDM. Fluorescence is reported as the normalized ratio or intensities at 350 nm and 330 nm.

### PSM mutants in each chain have different effects on the TmrAB heterodimer ATPase activity

TmrAB is derived from a thermophilic bacteria that exhibits optimal growth and function at 68°C [33]. To determine the effect of PSM mutations on catalysis in TmrAB at its physiological temperature, we employed a colorimetric endpoint assay using ascorbic acid to measure the rate of ATP hydrolysis (**Figure 5A; Table 1**). Because TmrAB contains a single functional NBD only in TmrA and the second site likely has ATP bound through the catalytic cycle, all data were fit with a single site Michaelis Menten model. WT TmrAB displayed characteristic ATPase activity with a measured *K*_*m*_ value of 1.35 ± 0.24 mM and a *k*_*cat*_ of 3.03 ± 0.19 s^−1^, consistent with previously published reports [10,33].

**Table 1.**
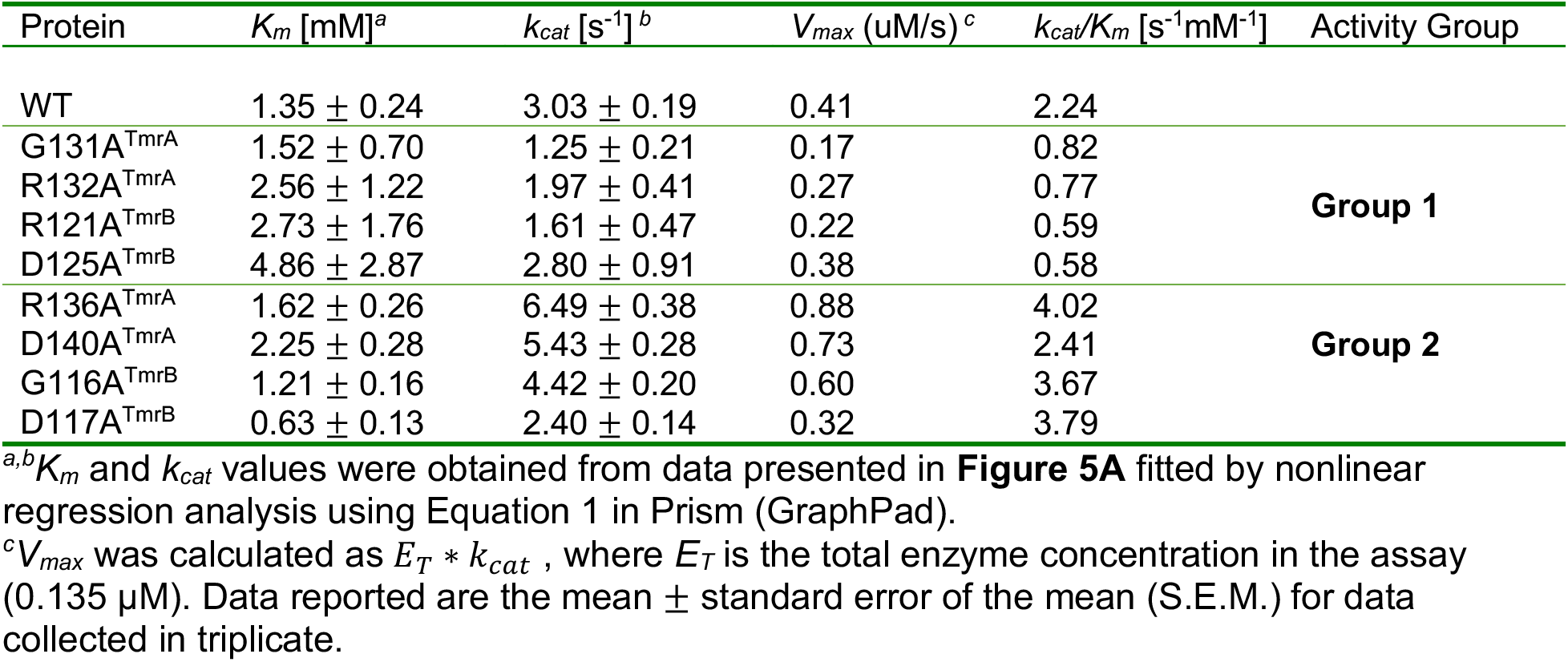
ATPase activity of TmrAB Peptide Sensor Motif mutants

**Figure 5.**
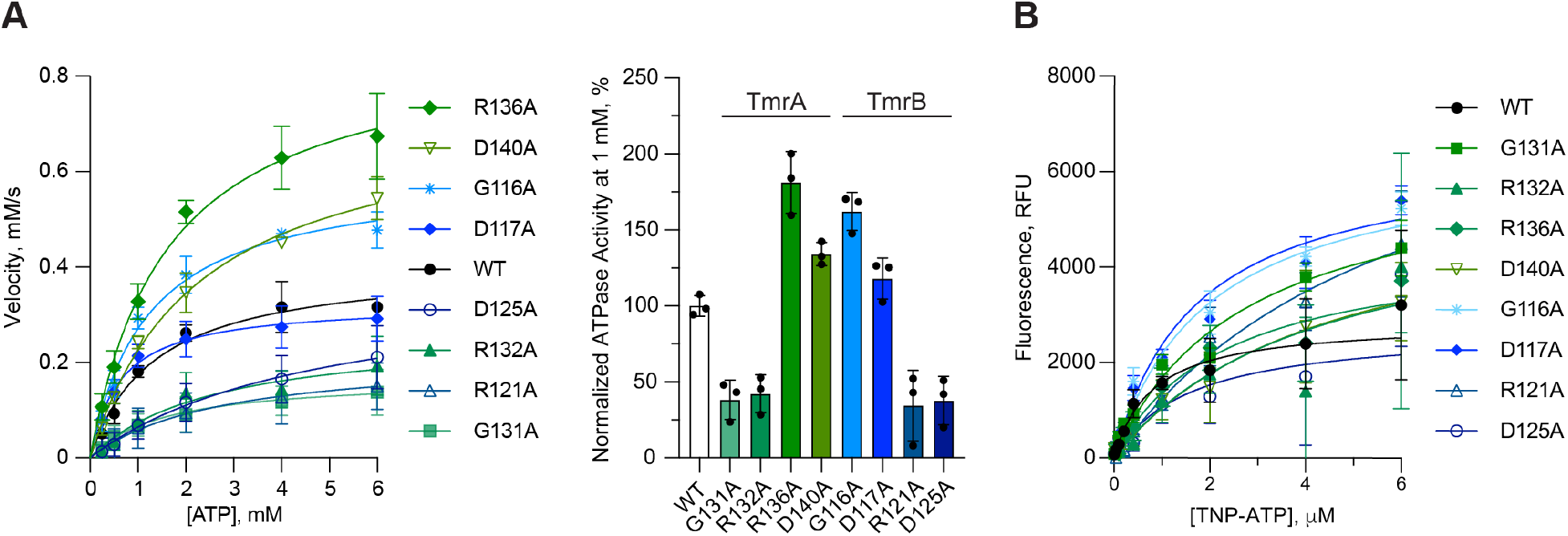
PSM mutations alter ATP hydrolysis rates differently in each chain of TmrAB independently of ATP binding. **A**) ATP hydrolysis activity at 68°C measured by ascorbic acid colorimetric assay for 0.135 μM WT or mutant TmrAB, and varying concentrations of ATP (*left*). ATPase rates in the presence of 1 mM ATP in this data are normalized to the mean rate for WT TmrAB (*right*). Michaelis-Menten kinetic parameters *K*_m_, *k*_cat_, and *k*_cat_/*K*_m_ were determined using Prism (GraphPad) and are summarized in **Table 1. B**) Binding curves generated from the change in fluorescence observed at room temperature for WT or mutant TmrAB (0.17 μM) incubated with increasing concentrations of the fluorescent ATP analog, TNP-ATP (Sigma). Data presented in the graphs in (****A****) and (****B****) are the mean ± standard deviation (S.D.) for four replicates (n=4). Normalized data reported in (****A****) are the mean ± standard error of the mean (S.E.M.). Apparent binding affinities for TNP-ATP (*K*_*D-app*_^*TNP-ATP*^) were determined using Prism (GraphPad) and are summarized in **Table 2**.

**Table 2.** Apparent nucleotide binding affinities for TmrAB Peptide Sensor Motif mutants

| Protein | $K_{D-app}^{TNP} [\mu M]^d$ | P-value<sup>e</sup> |
| --- | --- | --- |
| WT | $0.81 \pm 0.31$ | --- |
| G131A<sup>TmrA</sup> | $2.72 \pm 0.82$ | 0.07 |
| R132A<sup>TmrA</sup> | $4.26 \pm 4.09$ | 0.43 |
| R136A<sup>TmrA</sup> | $2.31 \pm 1.00$ | 0.20 |
| D140A<sup>TmrA</sup> | $4.50 \pm 1.32$ | 0.03 |
| G116A<sup>TmrB</sup> | $1.97 \pm 0.50$ | 0.10 |
| D117A<sup>TmrB</sup> | $1.72 \pm 0.36$ | 0.11 |
| R121A<sup>TmrB</sup> | $6.17 \pm 2.59$ | 0.09 |
| D125A<sup>TmrB</sup> | $1.57 \pm 1.52$ | 0.64 |

<sup>d</sup> $K_{D-app}^{TNP-ATP}$ values were obtained from data presented in **Figure 5B** fitted by nonlinear regression analysis using Equation 2 in Prism (GraphPad). Data reported are the mean ± S.E.M.
<sup>e</sup>P-value denotes probability of the null-hypothesis that the mutant mean is equal to that of WT TmrAB.

Changes in mutant activity can be divided into two groups (in each case compared to WT) as judged by k_*cat*_/K_*m*_, which was chosen because it provides an indirect measure of the ability of each productive ATP-bound complex to couple an ATP hydrolysis event to a conformational change, and thus informs on the effect of PSM mutations in conformational coupling (**Figure 5A, Table 1**). In group 1,catalytic efficiency was decreased to 26 – 37% of WT TmrAB in the G131A^TmrA^, R132A^TmrA^, R121A^TmrB^, and D125A^TmrB^ mutants. Interestingly, this effect was nearly inverted in group 2 which contained residues from the opposite chain as group 1. In group 2, activity increased ~1.7-fold in G116A^TmrB,^ D117A^TmrB^, and in R136A^TmrA^. Activity was only marginally increased (107% of WT) in D140A^TmrA^.

### PSM mutants do not influence ATP binding

To determine whether the effects on ATP hydrolysis for our PSM mutants arise from deficiencies in nucleotide binding affinity, we determined the apparent affinity, K_*D-app*_, for the fluorescent ATP analog, TNP-ATP, in WT TmrAB and each PSM mutant (**Figure 5B, Table 2**). The K_*D-app*_^*TNP-ATP*^ ranged from 0.81 μM in WT and 1.57 – 6.17 μM in PSM mutants. These values for the dissociation constant are significantly lower than their enzymatic K_*m*_ values, and likely due to formation of partially occluded states as is the case in PgP [20], but overall still consistent with previous reports of the K_*D-app*_^*TNP-ATP*^ in MsbA [39]. It is important to note that these K_D_ values are apparent K_D_ values since there are two nucleotide binding sites and it has been proposed that one site can be more occluded [6] and that this site has higher affinity [20]. There was no statistically significant difference in affinities for ATP between WT TmrAB and the PSM mutants, suggesting that the PSM residues are likely not a determinant for ATP binding.

### ATPase-deficient PSM mutations block entry into low peptide affinity conformations of TmrAB

According to the processive-clamp/switch model [16,18], transport is only achieved when the high affinity inward-facing state changes to a low affinity outward-facing state to release substrate. To test the ability of TmrAB PSM mutants to induce the IF-to-OF switch and thus enable substrate release, we employed a fluorescent polarization binding assay to quantify binding of a fluorescently-labeled peptide substrate (RRY(C^Fluorescein^)KSTEL) to TmrAB in the absence and presence of ATP (**Figure 6**). In this assay, a decrease in the polarization signal in response to ATP binding corresponds to a loss of bound peptide substrate, presumably through the ability of TmrAB to sample the low-affinity outward open conformation [6]. Our experiments were performed at 4°C where ATP hydrolysis is not observed in TmrAB to prevent the effects of trans-inhibition by hydrolysis products, and for concentrations of peptide where at least 30% of the total enzyme in the sample is bound to minimize protein consumption, under similar concentrations and conditions as previously described [10,31,40].

**Figure 6.**
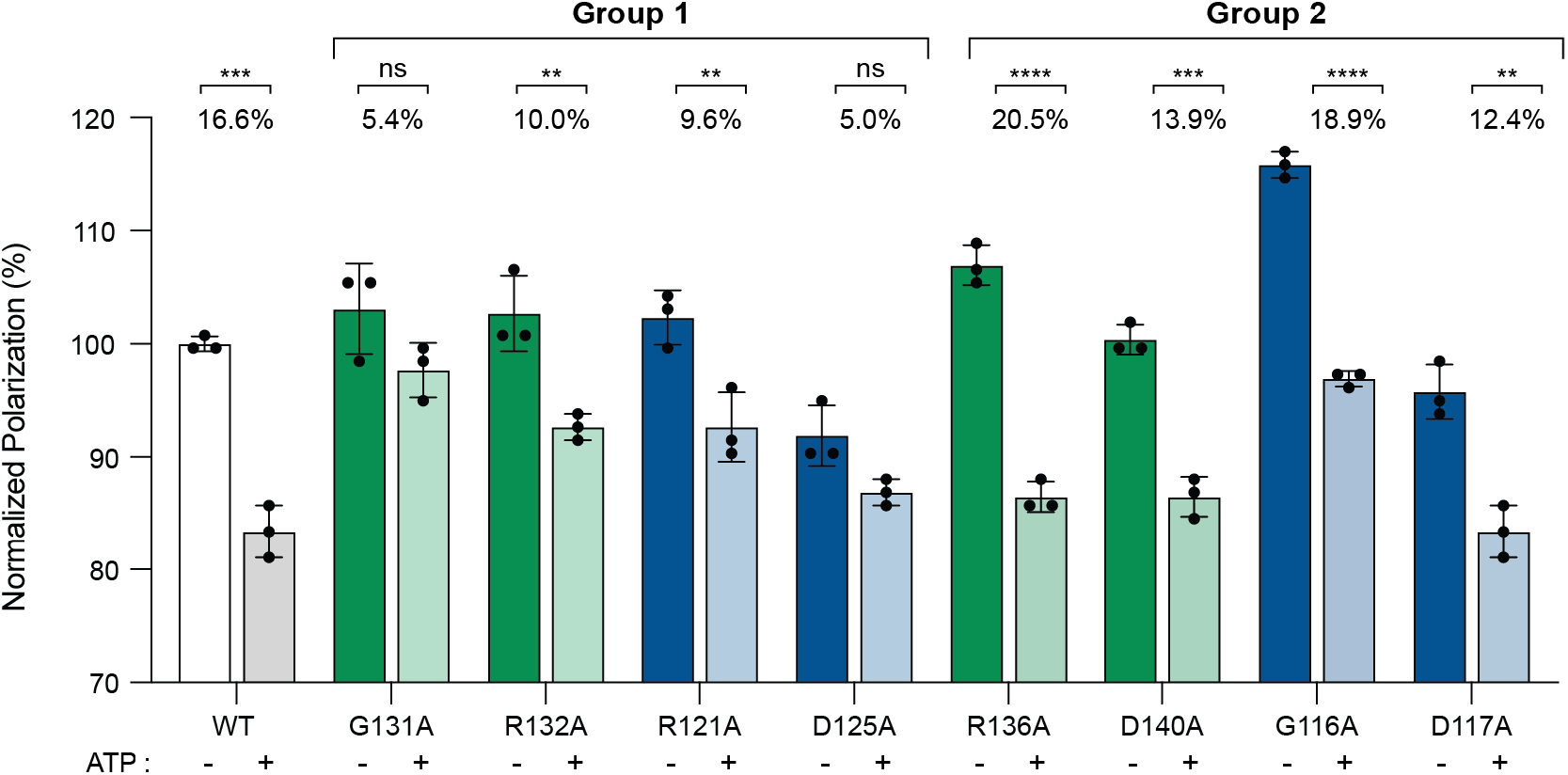
Peptide substrate release is affected in TmrAB mutants G131A^TmrA^ and D125A^TmrB^. The change in fluorescence polarization was monitored for WT and mutant TmrAB (1.7 μM) incubated with 50 nM of the fluorescent transport substrate peptide, RRY(C^Fluorescein^)KSTEL and in the absence (*green* bars in TmrA mutants; *blue* bars in TmrB mutants) or presence (*light green* bars in TmrA mutants; *light blue* bars in TmrB mutants) of 5 mM ATP added to achieve the outward-facing state for mutants in group 1 (decreased ATPase activity) and group 2 (unchanged or increased ATPase activity). Data are presented as the percentage mean ± S.D. for three replicates (n=3) normalized to the mean value for WT TmrAB in the absence of ATP. P-values are reported above the data (ns: not significant (P ≥ 0.05), **: P ≤ 0.01, ***: P ≤ 0.001, ****: P ≤ 0.0001).

The peptide binding data showed a decrease in peptide binding in the presence of ATP for WT TmrAB, as is expected. Group 1 mutants R132A^TmrA^ and R121A^TmrB^ and all group 2 mutants (R136A^TmrA^, D140A^TmrA^, G116A^TmrB^ and D117A^TmrB^) similarly showed this change, suggesting that they also can attain a low affinity state. These results suggest conformational coupling between the NBD and TMD proceeds as expected in these mutants in which ATP hydrolysis was increased. The change in peptide binding in the G131A^TmrA^ and D125A^TmrB^ variants was not significant, consistent with their decreased efficiency of ATPase activity. While peptide binding was also slightly decreased in an ATP-dependent manner in the ATPase deficient R132A^TmrA^ and R121A^TmrB^ variants, the observed change was less significant than in the catalytically-competent PSM mutants. Together, this data suggests that ATPase deficient PSM mutants (group 1) trapped in the high-affinity conformational state of TmrAB may not induce the IF-to-OF switch in the presence of ATP binding.

### Mutation of the 100% conserved PSM glycines block the ability of TmrAB to achieve an outward-facing conformation

To understand how the positioning of the PSM might influence the transport cycle, we focused on mutants of the two most conserved residues with opposite functional effects on catalytic activity and peptide binding, G131A^TmrA^ and G116A^TmrB^. In order to measure formation of exclusively the OFopen state of TmrAB, we engineered cysteines into the glycine variants for disulfide cross-linking on the outside gate of TmrAB. These double cysteine mutations correspond to positions previously investigated in the homodimeric transporter MsbA [39] (L290C in TmrA and V275C in TmrB) (**Figure 7A**), and are positioned such that a cross-link only forms in IFopen or OFocc states and cannot form in the OFopen state necessary for transport, thus informing on the formation of this state.

**Figure 7.**
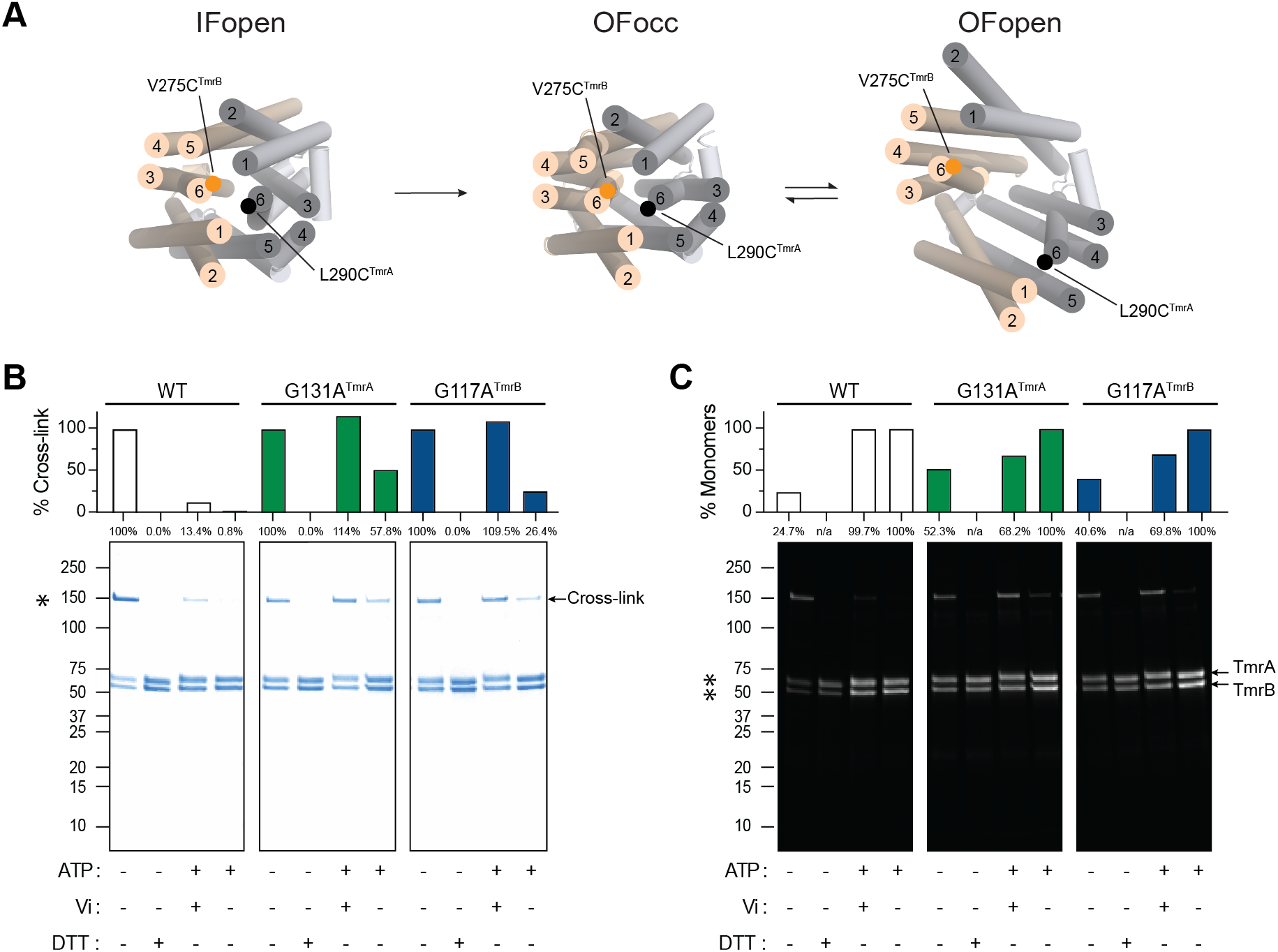
PSM mutations disrupt the formation of the outward-facing open conformation in TmrAB. **A**) Top view of relative transmembrane domain helix orientations in the IFopen (left, PDBID: 6raf [10]), OFocc (middle, PDBID: 6rak [10]), and OFopen (right PDBID: 6raj [10]) states of TmrAB highlighting the positions of double cysteine mutations introduced in the TmrA (L290C^TmrA^) and TmrB (V275C^TmrB^) chains of cysteine-less (C416S^TmrA^) WT TmrAB and G131A^TmrA^ and G116A^TmrB^ mutants. **B**) SDS-PAGE analysis of protein crosslinking in double cysteine mutants of WT and G131A^TmrA^ or G116A^TmrB^ TmrAB following incubation with and without ATP and/or sodium orthovanadate (Vi) at 68°C. The higher molecular weight bands (150 kDa) corresponding to the cross-linked TmrAB dimer were quantified for each condition and reported as a percentage relative to the amount of crosslinked TmrAB observed without substrate or DTT incubation. **C**) UV-illuminated image of the gel reported in (**B**) following treatment with ATTO-590 to detect free cysteines in monomeric TmrA and TmrB. The monomeric population in each condition was quantified and is reported the same as in (**B**). Samples treated with DTT were excluded from the ATTO-590 quantification (n/a) in (**C**).

In the absence of ATP, the majority of WT TmrAB forms the IFopen state as assessed by the predominant presence of a ~150kDa band on an SDS-PAGE gel corresponding to a dimer between TmrA and TmrB (**Figure 7B,C**). This result is consistent with similar previously performed studies on ABC exporters [39]. This higher molecular weight crosslinked band disappeared upon incubation with either ATP alone or ATP plus sodium orthovanadate (ATP-Vi), which traps an OF state as expected. However, when repeated with the PSM glycine mutants, our experiment showed an almost complete recovery of crosslinks in the ATP-Vi condition. This implies that the ATP-Vi bound state is not able to hold TmrAB in the OFopen conformation. Notably, a less pronounced effect was observed in the absence of sodium orthovanadate (ATP alone), with only a partial recovery of cross-linking. It is unclear why ATP-Vi or ATP alone have different effects in the mutants, though the cryo-EM structures of TmrAB find that datasets in the presence of vanadate have a particle distribution of ~65% and ~35% in the OFocc and OFopen respectively, whereas ATP alone primarily adopts the OFopen conformation [10]. Nevertheless, our cross-linking results suggest that both the G131A^TmrA^ variant and the G116A^TmrB^ have an impaired ability to couple the presence of ATP or ATP-Vi to the formation of OFopen.

## DISCUSSION

This work identifies the peptide sensor motif as part of an evolutionarily and functionally important latch located in a region of ICL-1 connecting the NBDs and TMDs. The latch connects two conserved electrostatic clusters at the start and end of the PSM which previously published cryo-EM structures show are formed only after transport is initiated and a gross conformational change is adopted. Our crosslinking results together with ATPase data on peptide sensor mutants show that PSM-mediated formation of the electrostatic latch is integral for both proper execution of the ATPase cycle and stabilizing the fully outward-open facing state. Changes to peptide binding in the presence of ATP in a subset of mutants show that destabilization of the state likely arises from a failure to properly execute the IF-to-OF transition necessary for transport. Together, our data identify a mechanism for how an allosteric signal between distal substrate bindings sites may be communicated during transport in asymmetric ABC exporters.

Protein allostery is diversely defined and can be executed through global conformational changes and domain movements [41], local rearrangements of side chain networks [42], or by subtle alterations in protein dynamics independently of a conformational change [43]. Analysis of evolutionarily conserved networks is a powerful approach for inferring conserved mechanisms of allostery from sequence. Here, two complimentary evolutionary analysis methods were used to identify such networks in the understudied, but critical, TmrAB exporter ICL region: Evolutionary Trace [35] to identify conserved *single residues* important for function and Evolutionary Couplings [36,37] to identify conserved contacts between *pairs of residues.* Evolutionary couplings are particularly informative since they can infer multiple types of interactions – those important for function, those necessary for stabilizing the folded state of a protein, and those that stabilize functionally important alternate conformations. Both methods clearly highlighted the ICL-1 as a central hub for functionally important interactions with several conserved sites clustered in the NBDs and TMDs. Unexpectedly, the most significant of these contacts form networks within one chain of TmrAB that largely do not cross-over into the other chain. However, one prominent network emerged from our analysis in which conserved residues of the canonical TmrA chain PSM act as a conduit bridging sites on the non-canonical TmrB NBD to the TmrA TMD. Notably this includes residues from the TmrB X-loop, which is overall more conserved than the X-loop in TmrA. Furthermore, nearly every pair predicted by evolutionary couplings could be observed forming a direct interaction when mapped onto recently determined cryo-EM structures of TmrAB in multiple conformational states including the inward-facing open state, which is the resting state of exporter type ABC transporters. Our melting data showed no discernable change in the stability of protein fold in the IFopen state in the presence of PSM mutations. Therefore, we conclude that these contacts formed upon global conformational changes, including NBD dimerization and the IF-to-OF transition, are primarily integral to the allostery between the NBD and TMD. Although other networks were not identified in this study, they cannot be ruled out until additional detailed molecular dynamics studies are conducted to test their possibility.

Interestingly, despite both halves of TmrAB undergoing somewhat similar conformational changes upon activation, every PSM mutant impairs TmrAB catalysis in a directly opposite ways from its analogous residue in the other chain of the heterodimer. It is unclear why such functional asymmetry in our data exists. PSM-mediated defects fall into two defined groups characterized by the effect of mutation on ATPase activity and peptide substrate binding. (**Table 3**). In group 1 (G131A^TmrA^, R132A^TmrA^, R121A^TmrB^ and D125A^TmrB^), the decreased ATPase activity relative to WT TmrAB and impaired formation of a low affinity substrate binding site in the presence of ATP suggests hydrolysis is rate-limited by the stability of the OFopen conformation. In contrast, the second group of mutants (G116A^TmrB^, D117A^TmrB^, R136A^TmrA^ and D140A^TmrA^) were marked by elevated ATPase activity compared to WT TmrAB suggesting a futile ATPase cycle as is observed for mutations in the Q-loop and X-loop of other ABC transporters [28,29]. Such futile cycles often uncouple ATPase activity from the transport cycle, which we also observe in TmrAB. Interestingly, very few of the evolutionarily important contacts, especially in the TMD, identified in our bioinformatics analyses are formed between TmrA and TmrB and instead are *intra*-chain. *Inter*-chain interactions were limited to those formed between the NBDs and ICLs (**Figure 2**), and the observed enzymatic asymmetry of the PSM mutants likely reflects the functional asymmetry of the NBD to which they are networked.

**Table 3.**
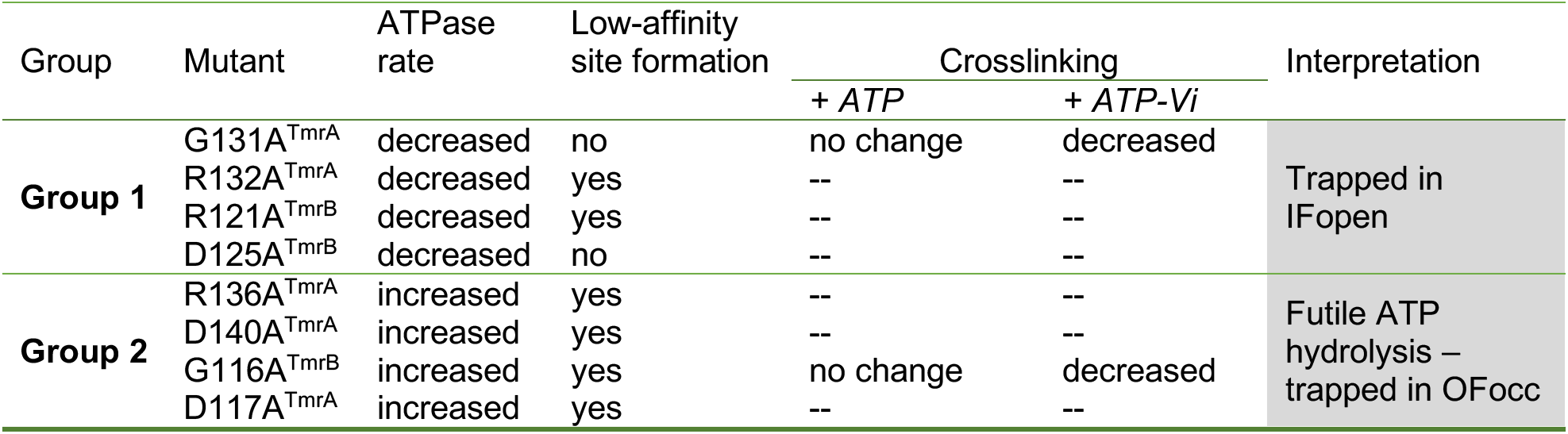
Summary of Peptide Sensor Motif mutation effects on TmrAB

| Group | Mutant | ATPase rate | Low-affinity site formation | Crosslinking |  | Interpretation |
| --- | --- | --- | --- | --- | --- | --- |
|  |  |  |  | + *ATP* | + *ATP-Vi* |  |
| **Group 1** | G131A<sup>TmrA</sup> | decreased | no | no change | decreased | Trapped in IFopen |
|  | R132A<sup>TmrA</sup> | decreased | yes | -- | -- |  |
|  | R121A<sup>TmrB</sup> | decreased | yes | -- | -- |  |
|  | D125A<sup>TmrB</sup> | decreased | no | -- | -- |  |
| **Group 2** | R136A<sup>TmrA</sup> | increased | yes | -- | -- | Futile ATP hydrolysis – trapped in OFocc |
|  | D140A<sup>TmrA</sup> | increased | yes | -- | -- |  |
|  | G116A<sup>TmrB</sup> | increased | yes | no change | decreased |  |
|  | D117A<sup>TmrA</sup> | increased | yes | -- | -- |  |

An unusual aspect of our data is the similarities in the functional properties of the PSM glycine mutants to those of the charged residues of the PSM in the opposite chain. The IF-to-OF switch may be aided by a cluster of glutamates that interact with the positive helix dipole of the conserved motif (E218^TmrA^ and E470^TmrB^ with the TmrA PSM; E203^TmrB^ and E492^TmrA^ with the TmrB PSM). Notably, E218^TmrA^ and E470^TmrB^ were ranked in the top 5% of functionally important residues predicted by ET analysis and are part of the continuous network connecting the Q-loop and X-loop of the TmrB NBD to the TmrA PSM and TMD substrate binding sites (**Figure 2B**). The stability imparted by these interactions may be the key element helping to propagate an allosteric signal from the NBD to the TMD.

The IF-to-OF transition does not result in a large change in the PSM backbone atom dihedral angles suggesting the flexibility of the nearly absolutely conserved glycines (G131^TmrA^ and G116A^TmrB^) is not important for PSM function. Instead, it is likely that the small size of glycine is necessary for both TM3^E^ helices harboring the PSMs to move past other parts of the intracellular loop region that rearrange and become tightly packed in the OF states of ABC exporters like TmrAB. This constricted assembly may be necessary to stabilize the non-typical π-helix geometry formed by residues of the PSM [44]. Frequently observed in critical conformational junctions of dynamic integral membrane proteins, π-helices are often formed and broken between structural transitions and are observed in the TM3^E^ helices of several transporters such as the bacterial amino acid transporter LeuT [44,45], in the OF states of ABC transporters [8,10,46], and in the human temperature sensitive ion channels of the TRPV family [47–49].

It is unclear if our results are specific to heterodimers or more broadly reflect homodimeric mechanisms in exporters that share the TmrAB fold (families B,C, and D in humans), though there is evidence that asymmetry may also be relevant for exporters with two canonical NBDs. Most significantly, asymmetry in ATP binding has been observed in the human drug transporter PgP, which has two functional NBDs [20]. Investigation of DARPin binding to the bacterial homodimer MsbA also showed asymmetric binding of a single DARPin at a time depending on the specific state of ATP along the hydrolysis cycle in each NBD [50]. Lastly, EPR distance measurements on PgP along its transport cycle reveal inherent conformational asymmetry similar to that observed for heterodimeric transporters, thus reflecting processive hydrolysis of ATP at two NBDs [14]. A later EPR study on PgP showed that asymmetric conformations are associated with the substrate bound states, but not for inhibitor bound states [51]. These proteins contain the same PSM-mediated conserved networks identified in our evolutionary analyses of TmrA and TmrB. This evolutionary significance supports a role of the PSM as a common feature of exporter type ABC transporters with the same fold as TmrAB independent of NBD functional symmetry.

In conclusion, we have identified the peptide sensor motif in TmrAB as an important element for the allosteric mechanism that couples ATP binding to conformational changes during transport. Mutation of the highly conserved motif not only results in alterations in catalysis, but also in an overall loss in the ability to attain an OFopen state. Findings from co-evolution analyses support a mechanism in which these effects likely arise from the destabilization of a conserved electrostatic network mediated by the PSM in TmrA. Specifically, we highlight a potential role of the almost absolutely conserved G131A^TmrA^ in aligning the NBDs and TMDs during the transition to the OF state. Together our results suggest the existence of differences in how conformational changes are propagated along each side of ABC exporters.

## MATERIALS AND METHODS

### Sequence alignments

Sequence alignments were performed against a collection of bacterial and mammalian ABC transporters that share the TmrAB fold. To limit hits to only ABC transporters with the TmrAB transmembrane fold, only the transmembrane sequence of TmrA (residues 1 – 332) and TmrB (residues 1-317) were used for the search. A blast search of the Uniref90 database for bacterial transporters was conducted with an E value cutoff of 0.01 to enable a robust search that covered wide evolutionary sequence space and multiple bacterial phyla. The sequences were then filtered to only include those containing a Walker B motif and that were between 540 and 900 amino acids (90% and 150% length of TmrA, respectively). To remove redundancy and downweigh closely related transporters, the sequences were next filtered through CD-HIT [52] with an 80% identity cutoff. Resultant sequences were aligned with default parameters in MAFFT [53,54] and manually inspected for gaps or undefined residues (1 sequence removed from TmrB alignments) resulting in 615 TmrA-like sequences and 567 TmrB-like sequences (**Supplemental Data 1**). Conservation of the PSM was analyzed using the WebLogo server [55]. The alignment presented in **Supplemental Figure 1** of a manual selection of representative ABC transporters from bacteria, yeast, and humans was also performed in MAFFT [53,54] [and colored by similarity in ESpript 3.0 [56]. A phylogenetic tree was generated on this alignment using raXML [57] and TidyTree (CDC).

### Evolutionary Trace analysis

Evolutionary trace analysis was performed using the custom MAFFT alignments described above. To facilitate analysis, the TmrA and TmrB sequences were respectively added back to the list of sequences that had been filtered for redundancy in CD-HIT [58]. The sequences were realigned with MAFFT [53,54] and uploaded to the UET server [36]. The rvET “coverage score” which ranks a given residue against all other residues in the protein was subtracted from 1 to generate a rank in which the most important residues were scored the highest. Residues ranked in the top 5% were mapped onto multiple structures of TmrAB to evaluate the relationship between functional importance and spatial proximity (**Supplemental Table 1**).

### Evolutionary couplings analysis

Evolutionary couplings in TmrAB were analyzed in three sets of alignments - TmrA (intra-TmrA – TmrA contacts), in TmrB (intra-TmrB-TmrB), and in TmrAB (inter-TmrA to TmrB) using the Evcouplings framework (**Supplemental Data 1**). To ensure sufficient coverage and robust coverage over as much of each TmrAB half as possible, JackHMMMR [59] was used to make a large sequence alignment of TmrA alone and TmrB alone with a Hidden Markov Model to fit difficult to align loops. HHsuite [60] was used to filter redundancy and cluster residues above an 80% sequence identity to ensure broad phylogenetic representation and to limit the overall number of sequences. Sequences were filtered using a Biopython [61] script for either a canonical or noncanonical Walker B motif to ensure proper comparison of either side of the heterodimer. With these criteria, 25,781 sequences were found related to TmrA and 5,000 sequences related to TmrB. Evcouplings analysis of the aligned sequences was then performed using PLMC and analyzed using tools from Evcouplings [37]. Comparison to known structures simultaneously carried out with Evcouplings.

### Molecular biology and cloning

Mutants of TmrAB were generated from a TmrAB template in the pet22b vector previously described in Zutz et. al. [33]. Mutagenesis primers were designed using the technique described by Liu and Naismith [62]. Constructs were verified by sequencing (Elim Biopharmaceuticals, Inc.). Construction of cysteine-less TmrAB templates (C416S) of WT TmrAB, G131A^TmrA^ and G116A^TmrA^ was performed by GenScript and used as the template for introducing two N-terminal cysteine residues at sites L290 and V275 in chains A and B, respectively.

### Protein expression and purification

WT and mutant TmrAB were expressed and purified using the method previously described by Zutz et. al [33]. Briefly, a starter culture of BL21(DE3) *E. coli* cells (NEB) transformed with TmrAB was incubated for 16 hours overnight with shaking at 37°C in 10 mL of Luria Broth (LB) supplemented with ampicillin. On the following day, 1L of Terrific Broth (TB) media was inoculated with 5 - 10 mL of the overnight culture and cells grown at 37°C until an OD_600_ of ~0.8 was reached. TmrAB expression was induced with 1 mM IPTG and cells cultured at 37°C for an additional four hours prior to harvest at 4°C by centrifugation at 5000xg for 20 min. For the G131A mutation, cells were cultured at room temperature for an additional 16 hours following induction.

Protein was purified first by resuspending cell pellets in lysis buffer (50 mM Tris-HCl, 300 mM NaCl, pH 8.0, and protease inhibitor tablets (Roche)) and lysed by pulse sonication on ice for 5 minutes in three cycles of 1 s on and 1 s off. Membranes were harvested from the cell lysate by ultracentrifugation at ~185,000xg for 1.5 hours at 4°C. The membrane pellets were then resuspended in buffer (50 mM Tris-HCl, 300 mM NaCl, 30 mM Imidazole, pH 8.0, and protease inhibitors (Roche)) at a ratio of 16 mL of buffer per 1 g of membrane pellet and solubilized to homogeneity by dounce homogenization. Then supplemented with iodoacetamide (1 mg/mL) except for the double cysteine mutants and ß-dodecyl maltoside detergent (ß-DDM, 1% w/v) for detergent extraction of membrane proteins overnight at 4°C with stirring. The clarified supernatant containing solubilized protein was collected by ultracentrifugation at ~150,000xg for 30 minutes prior to purification.

Detergent-solubilized TmrAB was next purified by loading onto a Nickel affinity column (HisTrap IMAC HP, GE Healthcare) and washing with binding buffer (50 mM Tris-HCl, 300 mM NaCl, pH 8.0, 0.1% ß-DDM) containing increasing concentrations of Imidazole (Wash 1: 30 mM Imidazole (10CV), Wash 2: 100 mM Imidazole (3CV)). Protein was eluted in binding buffer containing 300 mM Imidazole (2CV) and immediately diluted into Imidazole-free buffer and concentrated for size exclusion chromatography (SEC). SEC purification was performed at 4°C on an Enrich SEC 650 column (Bio-Rad) and sample eluted in SEC buffer (20 mM HEPES, 150 mM NaCl, pH 7, and 0.05% ß-DDM). Peak fractions containing pure TmrAB (WT or mutant) were pooled, concentrated, and stored at −80°C prior to further experiments.

### Intrinsic tryptophan and fluorescence analysis

To observe the effect of PSM mutations on TmrAB structural integrity and stability, changes in the emission spectra of tryptophan residues, induced by temperature-dependent unfolding, was monitored from 35°C to 95°C. TmrAB was diluted to 8.5 μM in SEC buffer and prepared in duplicate for measurement on a Tycho NT.6 (NanoTemper Technologies). Data are reported as the ratio of fluorescence intensity at 350 nm and 330 nm normalized to the minimum and maximum values of the whole dataset.

### Negative-stain analysis

Samples of WT and mutant TmrAB were prepared for negative-stain imaging to evaluate the quality and architecture of purified protein. Protein was diluted to ~0.2 μM concentration in SEC buffer and 3 μL applied to negative glow discharged carbon coated copper grids (EMS, Cat No. CF300-Cu-UL). Sample was incubated on grids for 30 s, then gently dipped in two drops of SEC buffer and one drop of 0.75% uranyl formate staining solution with blotting performed between drops by touching the grid edges to the surface of Whatman 1 filter paper. Grids were dipped in a second drop of stain for 30 s, blotted, and air dried prior to imaging. Images were collected on a 120 kV Tecnai Spirit (FEI) microscope equipped with an AMT XR80L-B 8 x 8 CCD camera at 49,000X magnification corresponding to a pixel size of 2.21 Å.

### Colorimetric ATPase assay

A colorimetric endpoint assay for inorganic phosphate determination described by Sarkadi et al. [63] was used to measure TmrAB ATPase activity near the physiological growth temperature of its host organism, *T. thermophilus* (68°C). Purified samples of WT or mutant TmrAB (1 to 10 μg) in 40 μL assay buffer (50 mM MOPS-Tris pH 7.0, 50 mM KCl, 150 mM NaCl, 0.05% ß-DDM and 2 μM DTT) were pre-incubated at 68°C for 5 minutes. The ATPase reaction was then started by the addition of 10 μL of Mg-ATP solution prepared in assay buffer, bringing the final concentration of TmrAB to 0.135 μM in a 50 μL reaction containing 10 mM MgCl_2_ and varying final concentrations of ATP (0-6 mM). Samples in the absence of Mg-ATP solution or TmrAB were prepared as negative controls. The reaction was quenched for various time points by adding 40 μL 5% SDS. Following quenching, 200 μL of detection reagent (8.75 mM Ammonium Molybdate, 3.75 mM Zinc Acetate, pH 5.0, and 7.5% Ascorbic Acid pH 5.0 prepared fresh prior to use) were added to each sample and incubated for 25 minutes at 37°C protected from light. The absorbance signal of total free phosphate in solution for each quenched sample and standards of KH_2_PO_4_ ranging in concentration from 0.0125 - 0.8 mM were subsequently measured at 800 nm in a clear 96-well plate (CoStar) on a a Synergy Neo2 Multi-Mode Microplate Reader (Biotek). Michaelis-Menten curves are shown as the mean ± standard deviation (S.D.) for data from three technical replicates (n=3) fit in Prism (GraphPad) with *Equation 1* below:

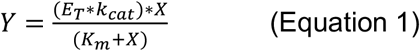

Where Y is the enzyme velocity (mM/s), E_T_ is the total enzyme concentration (mM), *k*_*cat*_ is the turnover number (1/s), X is the concentration of ATP (mM), and *K*_*m*_ is the Michaelis-Menten constant (mM). Kinetic parameters (*K*_*m*_, *k*_*cat*_, *V*_*max*_, and *k*_*cat*_/*K*_*m*_) are reported in **Table 1** as the mean ± standard error of the mean (S.E.M.).

### ATP binding assay

The binding affinity for nucleotide was measured as described in Doshi et. al. 2013 [39] and the apparent *K*_*D*_ for the fluorescent analog of ATP, TNP-ATP (Molecular Probes), calculated for WT TmrAB and PSM mutants. Samples were prepared in a total volume of 50 μL containing 0.17 μM WT or mutant TmrAB in SEC buffer supplemented with 1 mM MgCl_2_ and increasing concentrations of TNP-ATP (0-10 μM). Fluorescence data were immediately collected following the addition of TNP-ATP at 25°C in black 96-well half-area flat bottom microplates (CoStar) on a Synergy Neo2 Multi-Mode Microplate Reader (Biotek) at excitation and emission wavelengths of 408/10 nm and 535/15 nm, respectively. After subtracting for background fluorescence from TNP-ATP alone, data were normalized to the maximum fluorescence achieved within each dataset. Binding curves are shown as the mean ± S.D. for at least three technical replicates of WT TmrAB and PSM mutants fit in Prism (GraphPad) with a one-site specific binding model using *Equation 2* below:

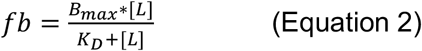

where *fb* is the fraction bound of protein with ligand, *B*_*max*_ is maximum binding, L is ligand, and K_*D*_ is the dissociation constant. Apparent dissociation constants for TNP-ATP binding (K_*D-app*_^*TNP-ATP*^) are reported in **Table 2** as the mean ± S.E.M.

### Peptide binding assay

Binding of the fluorescent transport substrate peptide RRY(C^Fluorescein^)KSTEL (Genscript) to WT and mutant TmrAB was determined as a function of the change in fluorescence polarization as described in [10]. In a total volume of 30 μL, 1.7 μM TmrAB in SEC buffer was incubated on ice for 15 minutes with 50 nM RRY(C^Fluorescein^)KSTEL and in the presence or absence of 5 mM ATP and 10 mM MgCl_2_. Fluorescence polarization data were collected at 25°C in black 96-well half-area flat bottom microplates (CoStar) on a Synergy Neo2 Multi-Mode Microplate Reader (Biotek) at excitation and emission wavelengths of 485/20 nm and 520/20 nm, respectively. Data are reported as the percentage mean ± S.D. for three replicates (n=3) normalized to the mean value for WT TmrAB in the absence of ATP.

### Cysteine cross-linking analysis

The ability of TmrAB to reach the OFopen conformation was assessed by cysteine disulfide cross-linking formation for TmrAB mutants in the presence or absence of ATP. A 1 μM solution of the C416S^TmrA^/L290C^TmrA^/V275C^TmrB^ mutants of WT TmrAB, G131A^TmrA^, and G116A^TmrB^ in SEC buffer was incubated with Mg-ATP solution (5 mM ATP, 10 mM MgCl_2_) plus 8 mM sodium orthovanadate for 3 min at 68°C, followed by the addition of 20 μM ATTO590-maleimide (Atto-tec) and incubating at room temperature for an additional 5 min. Subsequently, disulfide cross-linking reactions were initiated by the addition of 0.5 mM copper phenanthroline solution prepared as previously described in [39]. Samples were then incubated again at 68°C for an additional 15 min. Reactions were stopped by the addition of 10 mM N-ethylmaleimide (added from a freshly prepared stock of 100 mM prepared in ultrapure H_2_O) and incubation at room temperature for 2 min. A 10 μL aliquot from each reaction was electrophoresed on 4-20% polyacrylamide gels (BioRad), then viewed under UV light (at excitation and emission wavelengths of 605/650 nm) prior to Coomassie Blue-staining for visualizing all protein bands.

## ABBREVIATIONS

ABC transporter: ATP-binding Cassette transporter
ATP: adenosine 5′-triphosphate
B_*max*_: maximum ligand binding
CH1: coupling helix 1
CH2: coupling helix 2
Cryo-EM: cryogenic electron microscopy
DTT: dithiothreitol
EM: electron microscopy
ICL-1: intracellular loop 1
ICL-2: intracellular loop 2
ICL: intracellular loop
IF-to-OF: inward-facing to outward-facing
IFopen: inward-facing open
IMAC: immobilized metal affinity chromatography
K_*cat*_: turnover number
K_D-app_: apparent dissociation constant
K_D_: dissociation constant
K_*m*_: Michaelis constant
NBD: nucleotide binding domain
NS-EM: negative-stain electron microscopy
OFocc: outward-facing occluded
OFopen: outward-facing open
PSM: peptide sensor motif
SEC: size exclusion chromatography
ß-DDM: n-Dodecyl-ß-D-Maltoside
TM: transmembrane
TMD: transmembrane domain
TNP-ATP: 2′,3′-O-(2,4,6-Trinitrophenyl) adenosine 5′-triphosphate
Tris-HCl: Tris(hydroxymethyl)aminomethane hydrochloride
WT: wild type

## ACKNOWLEDGEMENTS

We thank Nitesh Khandelwal in the Tomasiak lab and Wolfgang Peti for helpful discussions and critical reading of this manuscript. Negative-stain images were collected at the Life Sciences North Imaging Facility at the University of Arizona. We thank Olivier Lichtarge and Panos Katsonis for technical assistance with Evolutionary Trace methods, and Kelly Brock for assistance with Evolutionary Couplings analysis. This was work was supported by grants from the National Institute of General Medicine Sciences to T.M. Tomasiak (R00 GM11424).

## AUTHOR CONTRIBUTIONS

C.R.M. and M.F. expressed all TmrAB samples. C.R.M. purified all TmrAB samples and performed biochemical assays. T. M. Thaker designed and performed site-directed mutagenesis to generate TmrAB mutants and collected microscopy data. V.F.T. assisted with molecular biology. T.M. Tomasiak performed and analyzed the large-scale sequence alignments. T.M. Tomasiak, T. M. Thaker, and C.R.M. designed the experiments, processed experimental data, and wrote the manuscript. T.M. Thaker and T. M. Tomasiak interpreted experimental results. T. M. Tomasiak planned and coordinated the project.

## SUPPLEMENTAL MATERIALS

**Supplemental Data 1. Sequence alignments and evolutionary analysis of TmrA and TmrB.** Aligned sequences used for Evolutionary Trace and Evolutionary couplings analysis related to Figure 1B, Figure 2, and Figure 3. Data can be accessed from: http://dx.doi.org/10.17632/c95j95r6nw.1

**Supplemental Table 1.** Evolutionary Trace Mapping Parameters

| Query Sequence | Q72J05 (TmrA) | Q72J04 (TmrB) | Q72J05 (TmrA) | Q72J04 (TmrB) |
| --- | --- | --- | --- | --- |
| Alignment window | 1-600 (600 residues) – All | 1-578 (578 residues) – All | 1-600 (328 residues) – All | 1-578 (317 residues) – All |
| Alignment | Jackhmmer 3.3, Uniref90, 5 iterations | Jackhmmer 3.3, Uniref90, 5 iterations | MAFFT, Uniref90 | MAFFT, Uniref90 |
| Filtering | 90% gapped columns and rows removed, 80% identity with HHSuite | 90% gapped columns and rows removed, 80% identity with HHSuite | 80% identity with CD-HIT | 80% identify with CD-HIT |
| Filtering for canonical or noncanonical NBD | Biopython | Biopython | Biopython | Biopython |
| N | 25,781 sequences | 4262 sequences | 624 (+ TmrA) | 570 (+ TmrB) |
| Sequences/L | 42.97 | 7.37 | - | - |
| Coevolution algorithm | plmDCA (EV couplings) | plmDCA (EV couplings) | Evolutionary Trace | Evolutionary Trace |
| Significance cut-off | P(couple) 0.99 | P(couple) 0.99 | Top 5%, Top10% | Top 5%, Top 10% |
| Structures analyzed | TmrA (6raf, 6rag, 6rah, 65ai, 6raj, 6rak, 6ran, 6ram, 5mkk) | TmrB (6raf, 6rag, 6rah, 65ai, 6raj, 6rak, 6ran, 6ram, 5mkk) | N/A | N/A |

**Supplemental Figure 1.**
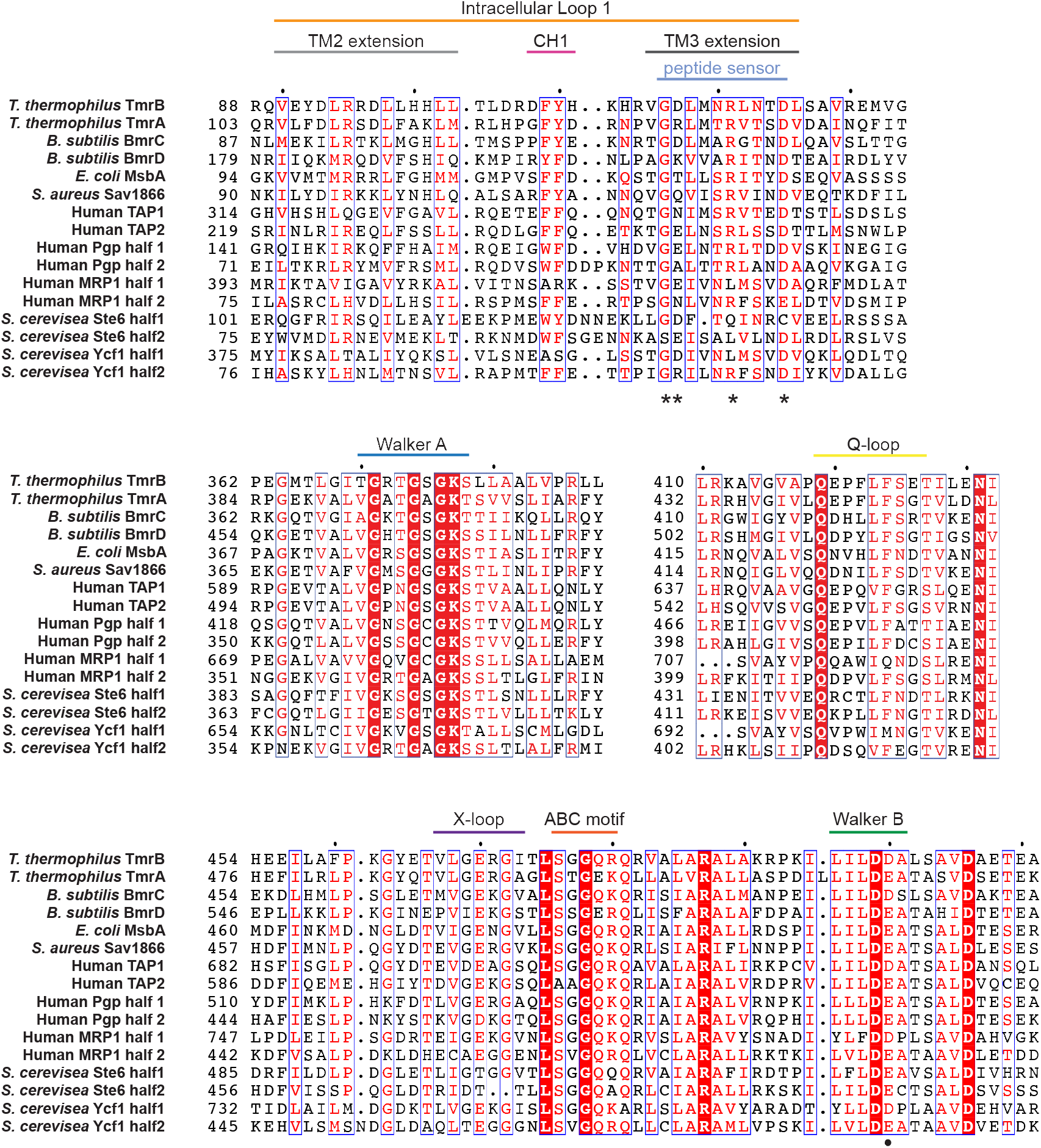
Sequence alignment of conserved elements in representative members of ABC transporter families. Sequences were aligned in MAFFT [52,53] and colored in ESPript 3.0 [64] by residue similarity. For the transporters that contain two nucleotide binding domains on a single polypeptide chain (PgP, MRP1, Ste6, and Ycf1) the sequence containing the second nucleotide binding domain (half 2) was included in the alignment as a separate sequence and aligned against the first half of the sequence containing the first nucleotide binding domain (half 1). Residues of the peptide sensor mutated in this study are denoted by an asterisk. The catalytic glutamate (E) in the Walker B motif is highlighted with a black circle below the sequence. Sequences containing an aspartate (D) in this position are considered non-functional for ATP hydrolysis.

**Supplemental Figure 2.**
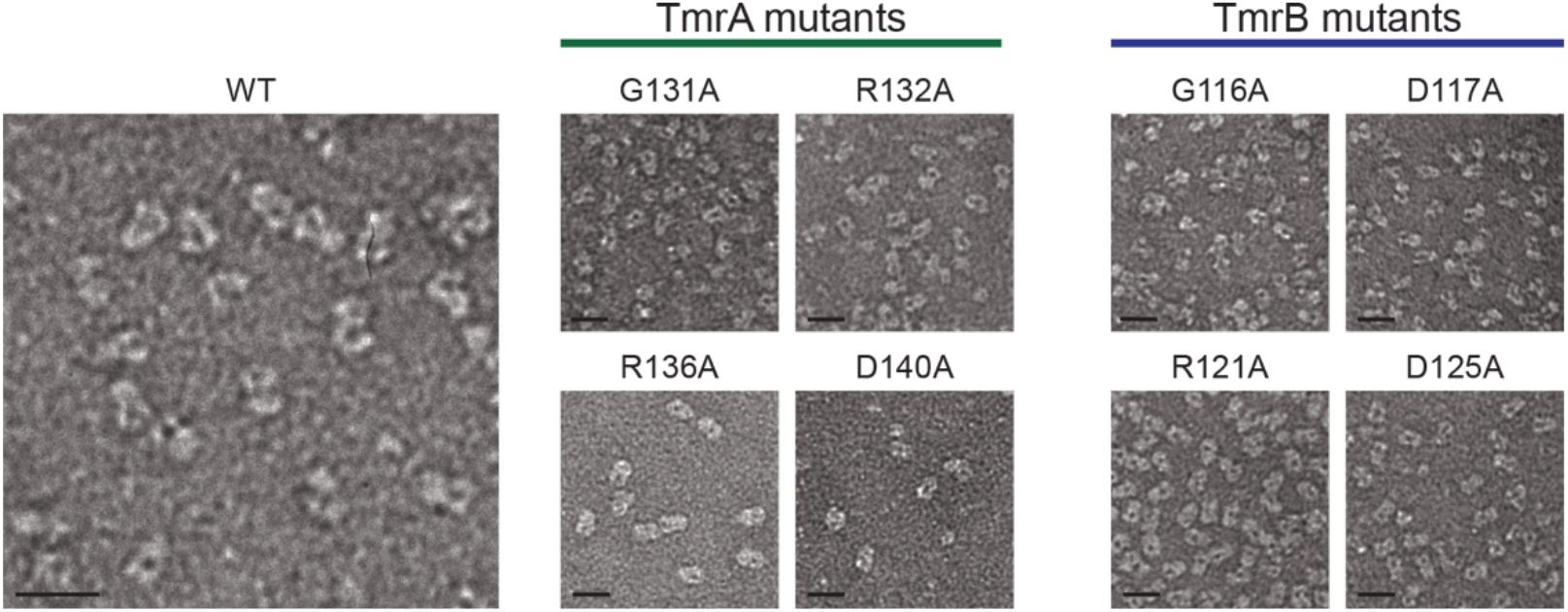
Negative-stain micrographs of purified wild-type (WT) and peptide sensor mutants of TmrAB collected at 49,000X magnification on a 120kV Tecnai Spirit microscope (FEI). Scale bars correspond to 20 nm.

